# Comparative analysis of histone H3K4me3 modifications between early embryos and somatic tissues in cattle

**DOI:** 10.1101/2020.11.11.379057

**Authors:** Mao Ishibashi, Shuntaro Ikeda, Naojiro Minami

## Abstract

Epigenetic changes induced in the early developmental stages by the surrounding environment can have not only short-term but also long-term consequences throughout life. This concept constitutes the “Developmental Origins of Health and Disease” (DOHaD) hypothesis and encompasses the possibility of controlling livestock health and diseases by epigenetic regulation during early development. As a preliminary step for examining changes of epigenetic modifications in early embryos and their long-lasting effects in fully differentiated somatic tissues, we aimed to obtain high-throughput genome-wide histone H3 lysine 4 trimethylation (H3K4me3) profiles of bovine early embryos and to compare these data with those from adult somatic tissues in order to extract common and typical features between these tissues in terms of H3K4me3 modifications. Bovine blastocysts were produced *in vitro* and subjected to chromatin immunoprecipitation-sequencing analysis of H3K4me3. Comparative analysis of the blastocyst-derived H3K4me3 profile with publicly available data from adult liver and muscle tissues revealed that the blastocyst profile could be used as a “sieve” to extract somatic tissue-specific modifications in genes closely related to tissue-specific functions. Furthermore, principal component analysis of the level of common modifications between blastocysts and somatic tissues in meat production-related and imprinted genes well characterized inter- and intra-tissue differences. The results of this study produced a referential genome-wide H3K4me3 profile of bovine early embryos and revealed its common and typical features in relation to the profiles of adult tissues.

**Supplementary information:** Supplementary data are submitted along with the main manuscript. The ChIP-seq datasets for bovine blastocysts have been deposited in the Gene Expression Omnibus of NCBI with accession number GSE161221.

## 1 Introduction

The periconceptional period of mammalian embryonic development is a critical window during which diverse environmental factors surrounding the embryo have not only short-term consequences such as effects on immediate embryonic development, but also long-term consequences including lasting influences on metabolic, developmental, and etiologic processes throughout gestation and even during postnatal and adult life (Sun, et al., 2015). During preimplantation development, dynamic epigenetic rearrangements occur, including substantial changes in DNA methylation and histone modifications, which regulate specific and heritable patterns of gene expression (Liu, et al., 2016; Wang, et al., 2018; Wang, et al., 2014). As the epigenome is dynamically formed during the preimplantation period, environmental intervention-induced changes in epigenome formation during this period have been considered as one of the possible causes of the long-lasting influences induced by the periconceptional environment (Sun, et al., 2015). This concept constitutes the “Developmental Origins of Health and Disease” (DOHaD) hypothesis and encompasses the possibility of controlling health and diseases in later life by epigenetic regulation during early development.

Some phenotypic changes in the field of livestock production can be discussed in the context of DOHaD given that early life events during prenatal and early postnatal development often affect traits, including those of economic importance (Chavatte-Palmer, et al., 2015; Gonzalez-Bulnes, et al., 2016; Sinclair, et al., 2016). For example, *in vitro* handling of ruminant early embryos in assisted reproductive technology increases the risk of fetal overgrowth syndrome (Chen, et al., 2015; Young, et al., 1998), whereas maternal nutrition, stress, or illness during pregnancy can affect productive traits such as postnatal growth, milk yield, carcass composition, and fertility (Chavatte-Palmer, et al., 2015; Gonzalez-Bulnes, et al., 2016; Sinclair, et al., 2016).

Although the epigenetic modifications that occur during the early developmental period, which persist in differentiated tissues, are considered a major mechanism of DOHaD, there is few comparative studies of histone modifications between early embryos and fully differentiated somatic tissues (Huang, et al., 2019). As a preliminary step for examining the changes of epigenetic modifications in early embryos and their long-lasting effects in later life, the elucidation of the common and typical features of epigenetic modifications between these two developmental stages is worth studying. In particular, accumulating evidence suggests that histone modifications at developmentally important genes, compared with DNA methylation, are more susceptible to the surrounding environment during the preimplantation period (Feuer, et al., 2014; Kudo, et al., 2015). In the present study, we aimed to obtain genome-wide profiles of histone H3 lysine 4 trimethylation (H3K4me3), a representative marker of active chromatin, in bovine blastocysts and to compare these profiles with those of adult liver and muscle tissues, which have been deposited in public databases. We examined whether we can extract the common and typical H3K4me3 features of early embryos and adult somatic tissues.

## 2 Materials and Methods

### 2.1 In vitro production of bovine embryos

This study was approved by the Animal Research Committee of Kyoto University (Permit Numbers 31-10) and was carried out in accordance with the Regulation on Animal Experimentation at Kyoto University. The bovine ovaries used in the study were purchased from a commercial abattoir as by-products of meat processing, and the frozen bull semen used for *in vitro* fertilization (IVF) was also commercially available. *In vitro* production of bovine embryos by IVF was performed as previously described (Ikeda, et al., 2018) except for that 50-μL drops of culture medium were used in *in vitro* culture after IVF. Blastocyst-stage embryos at 192 h post IVF were collected as approximately 11 embryos per biological replicate for Chromatin Immunoprecipitation (ChIP).

### 2.2 ChIP

ChIP for small cell numbers was performed with a Low Cell ChIP-Seq Kit (Active Motif) according to the manufacturer’s manual (version A3) with some modifications. The blastocysts were freed from the zona pellucida by using pronase before crosslinking with formaldehyde. After crosslink-quenching, the sample was sonicated to shear chromatin using a Bioruptor UCD-250 (Cosmo Bio) for 30 x 30 s with 30-s pauses in ice-water. The sample was centrifuged for 2 min at 18,000 x g and the supernatant (200 μL) was transferred to a new tube. The 200-μL sample of sheared chromatin was divided into a 10-μL aliquot as “input” and the rest (190 μL). The latter aliquot was processed for ChIP using 3 μg anti-H3K4me3 antibody (pAb-003-050, Diagenode) as indicated in the user manual. The input and ChIPed DNA was decrosslinked, purified, and resuspended in 40-μL low-EDTA TE buffer. The DNA samples were processed for library preparation for next-generation sequencing by using a Next Gen DNA Library Kit (Active Motif) following the manufacturer’s manual. The specific enrichment of H3K4me3 in the ChIP-seq libraries was validated by quantitative PCR for positive (1st exon-intron boundary of *GAPDH*) and negative (2nd exon of *MB*) regions.

### 2.3 Sequencing and data processing

Sequencing was performed on a HiSeq2500 (Illumina) as sigle-end 51-base reads. The sequencing reads were quality checked and aligned to the bovine genome (Bos_taurus_UMD_3.1.1/bosTau8, June 2014) except for scaffolds using Bowtie (Langmead, et al., 2009). The mapping duplicates were removed by Picard (http://broadinstitute.github.io/picard/). The peaks were called in the ChIP samples relative to the respective input samples using MACS (Zhang, et al., 2008). The annotation of called peaks to genomic regions was generated using CEAS (Shin, et al., 2009) and the peak occupancy rates were in its output. Average H3K4me3 enrichment profiles and heat maps were generated by ngs.plot (Shen, et al., 2014), and peak areas were calculated from its output. Gene ontology analysis was performed using the DAVID tool (Huang da, et al., 2009; Huang da, et al., 2009). ChIP-peaks were visualized using the Integrative Genomics Viewer (IGV) (Robinson, et al., 2011). The publicly available raw data for bovine liver and muscle were processed as described above except for the lack of input sample in the muscle sample. The common and specific peaks between samples were identified using bedtools (https://bedtools.readthedocs.io/en/latest/) with the default and -v option, respectively. Principal component analysis (PCA) was performed using SPSS with autoscaling of peak areas (SPSS Inc.).

### 2.4 Publicly available data

The following raw data from publicly available databases were used: ChIP-seq of bovine liver, Bull4 and Bull5 of E-MTAB-2633 (Villar, et al., 2015); ChIP-seq of bovine muscle (longissimus dorsi), GSM1517452 of GSE61936 (Zhao, et al., 2015); and RNA-seq of bovine blastocysts, GSM1265773, GSM1265774, and GSM1265773 of GSE52415 (Graf, et al., 2014). For RNA-seq data, the three datasets were merged and expression levels in RPKM values were calculated as previously described (Ishitani, et al., 2020). The genes were evenly divided into three categories as high, medium, and low expression levels according to the calculated RPKM values.

## 3. Results

### 3.1 H3K4me3 profile in bovine blastocysts

We performed ChIP-seq analysis of H3K4me3 using three biological replicates of bovine blastocysts (n = ~11 per replicate) derived from two independent IVF procedures. Pairwise comparisons of the ChIP signals in the biological replicates showed the high reproducibility of our method (Supplementary Fig. S1). We detected about 20,000 significant peaks throughout the genome (Supplementary Table S1 and Fig. 1A). Approximately 20% of the peaks were located on gene promotor regions (Fig. 1B). Figure 1C shows a snapshot of the H3K4me3 landscape in a 50-kb region (chromosome 5) that encompasses the transcription start site (TSS) of *GAPDH*, which is a representative positive region for H3K4me3 modifications (Herrmann, et al., 2013). Figure 1C demonstrates the clear enrichment of H3K4me3 at this region. Average profile plotting of the H3K4me3 signal around the genome-wide TSS regions showed similar profiles among the replicates and exhibited asymmetric bimodal peaks with a valley at TSSs that are considered to be nucleosome-free regions (Jiang and Pugh, 2009) (Fig. 1D). As expected, average profile plotting around gene bodies categorized by gene groups with different expression levels revealed that highly expressed gene groups had more extensive H3K4me3 modifications (Fig. 1E).

**Figure 1.**
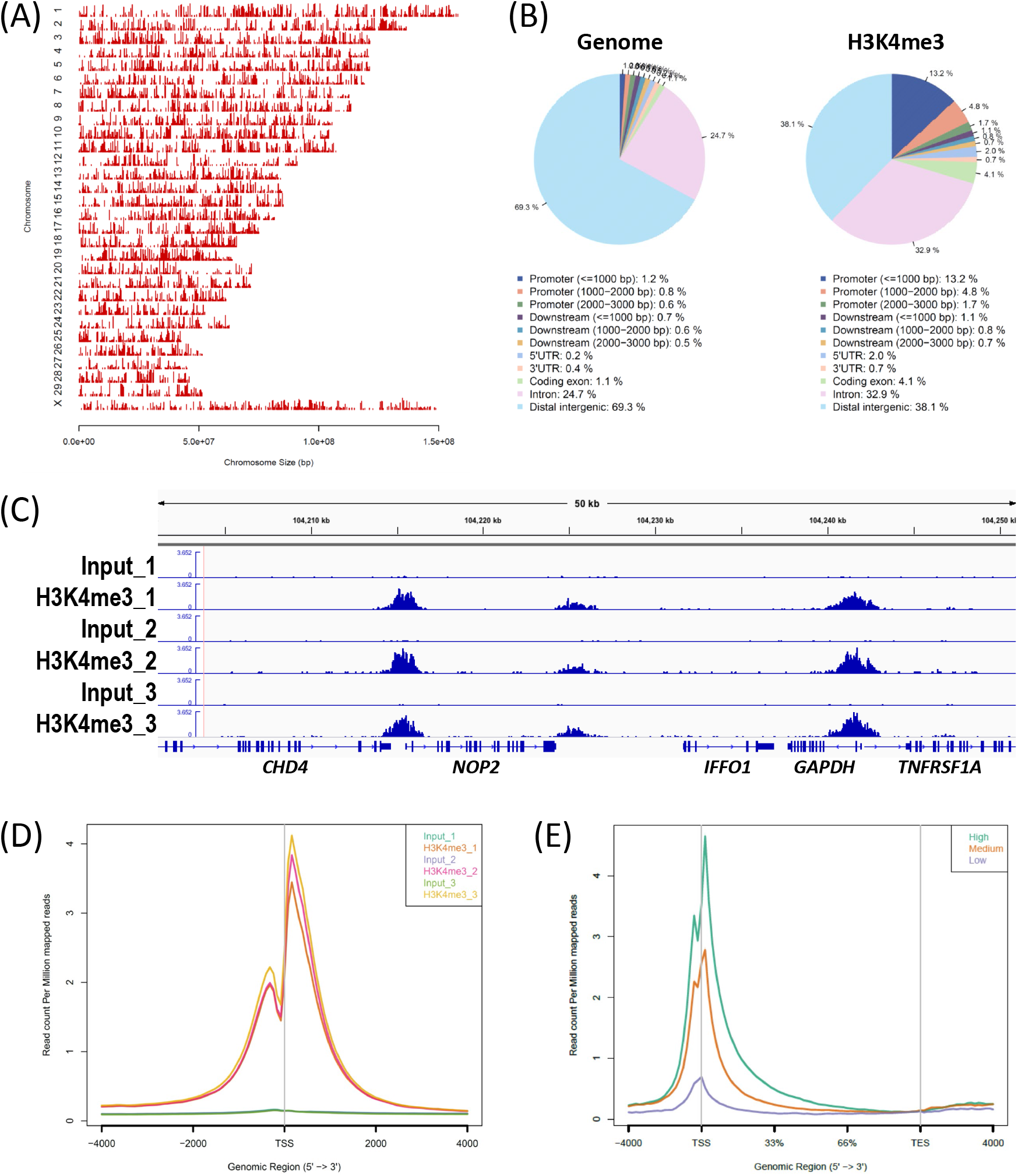
Overview of H3K4me3 ChIP-seq results in bovine blastocysts. (A and B) Distribution of H3K4me3 peaks in each chromosome (A) and in corresponding genic and intergenic regions (B). These figures were generated by the CEAS (Shin, et al., 2009) tool using the H3K4me3_1 (Blastocyst_1) sample. (C) Snapshot of the H3K4me3 landscape in a 50-kb region (chromosome 5) that encompasses the *GAPDH* TSS. The ChIP peaks in the three biological replicates were visualized using the Integrative Genomics Viewer (Robinson, et al., 2011). (D) Average profile plot of the H3K4me3 signal around the genome-wide TSSs. The three biological replicates are shown. (E) Average profile plot around gene bodies categorized by gene groups with different expression levels based on GSE52415 (Graf, et al., 2014). The H3K4me3_1 sample was used to generate the figure. The average profile plots were generated by ngs.plot (Shen, et al., 2014).

### 3.2 Characterization of embryonic- and somatic tissue-specific H3K4me3 modifications

We explored the tissue-specific H3K4me3 modifications between preimplantation embryos and somatic tissues. First, 14,018 overlapping peaks identified from two liver ChIP datasets in a public database (bulls 4 and 5 in E-MTAB-2633 (Villar, et al., 2015)) were designated as the liver peaks. On the other hand, the 20,298 overlapping peaks identified from our ChIP data from blastocysts (blastocysts 1 and 3, which exhibited the two highest peak numbers) were designated as the blastocyst peaks. Then, we merged these peak groups and extracted 1,899 and 7,901 liver- and blastocyst-specific peaks, respectively (Fig. 2A). From the genes that harbored these tissue-specific peaks within ±3,000 bp of the TSS, those with the top 100 peak occupancy rates were subjected to gene ontology (GO) analysis using the web-based DAVID tool (Huang da, et al., 2009; Huang da, et al., 2009). As a result, the genes with liver-specific peaks enriched the GO terms closely related to liver function such as “organic acid metabolic processes”, whereas the genes with blastocyst-specific peaks significantly enriched embryonic development-related GO terms such as “embryo development” and “cell fate commitment” (Fig. 2B). Such tissue function-related GO enrichment was not obtained from the liver and blastocyst peaks, without subtraction of the common peaks; in other words, analysis using these peak-associated genes generated only common and broad GO terms (Table S2). Figure 2C shows the representative liver- and blastocyst-specific peaks around the TSSs of *ARG1* and *GATA2*, respectively.

**Figure 2.**
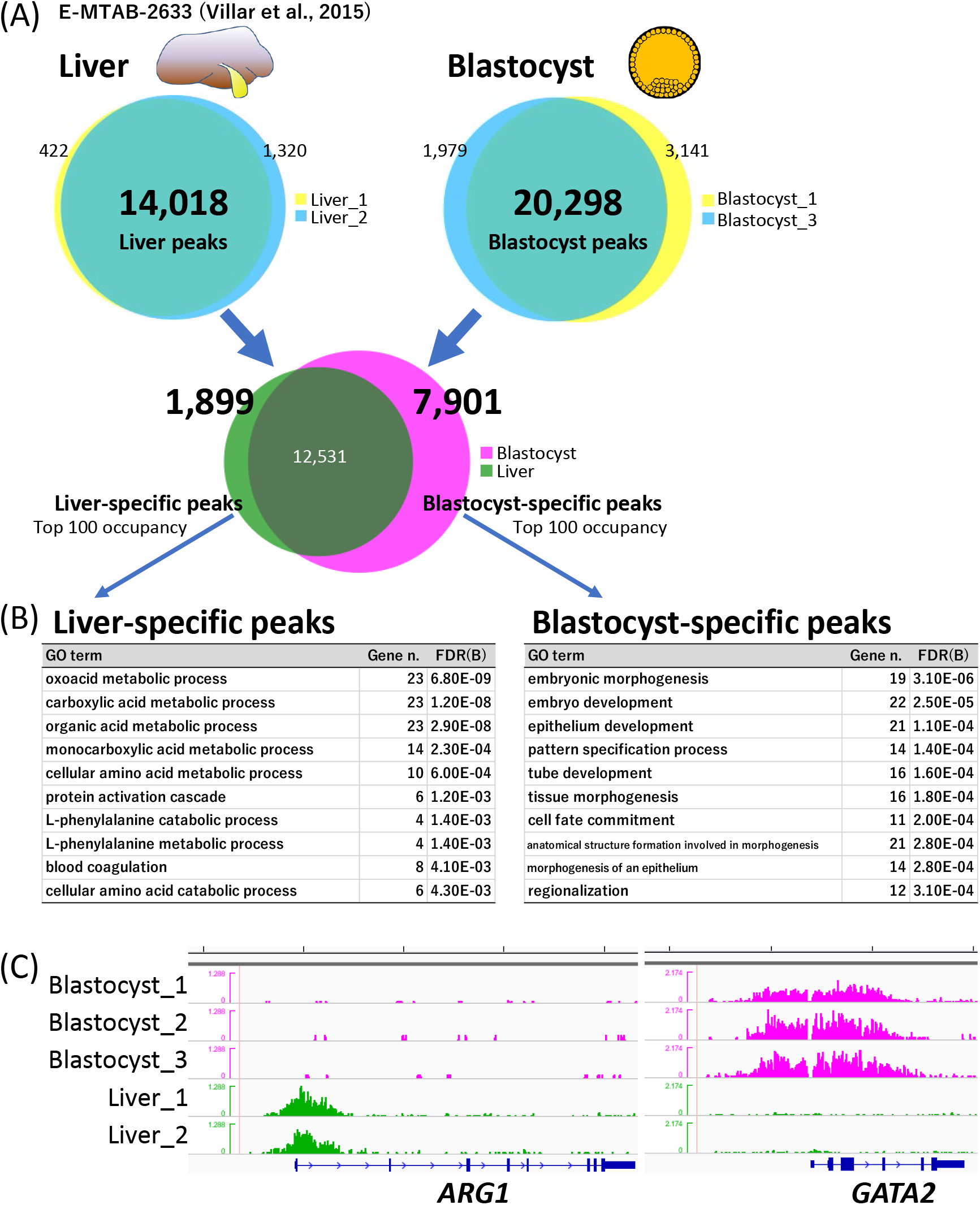
Characterization of blastocyst- and liver-specific H3K4me3 modifications. (A) Processing of ChIP-peaks in liver and blastocyst samples. The numbers show those of the peaks. The sum of common (intersect) and specific peak numbers is not equal to the original peak number in each sample because some peaks were separated into multiple peaks to represent intersects. (B) Top 10 significant GO terms for biological process enriched by the genes with the top 100 highest peak occupancy rates. Gene n. represents the numbers of related genes. FDR(B) indicates the Benjamin false discovery rate. (C) Examples of the H3K4me3 landscape of liver- (*ARG1*) and blastocyst-specific (*GATA2*) peaks around their TSSs. Five-kb graduations are shown in the top scale.

We applied the same strategy to characterize muscle tissue in terms of the H3K4me3 profile. We used publicly available H3K4me3 data from the longissimus dorsi muscle of beef cattle (Zhao, et al., 2015) and compared them with our blastocyst data. GO analysis using the muscle peaks generated broad terms again; however, the top 10 significant GO terms generated by the muscle-specific peaks extracted by comparison with the blastocyst peaks all contained the word “muscle” (Fig. 3A and B). The genes participating in the enriched GO terms identified muscle function-related genes with H3K4me3 modifications (Fig. 3C).

**Figure 3.**
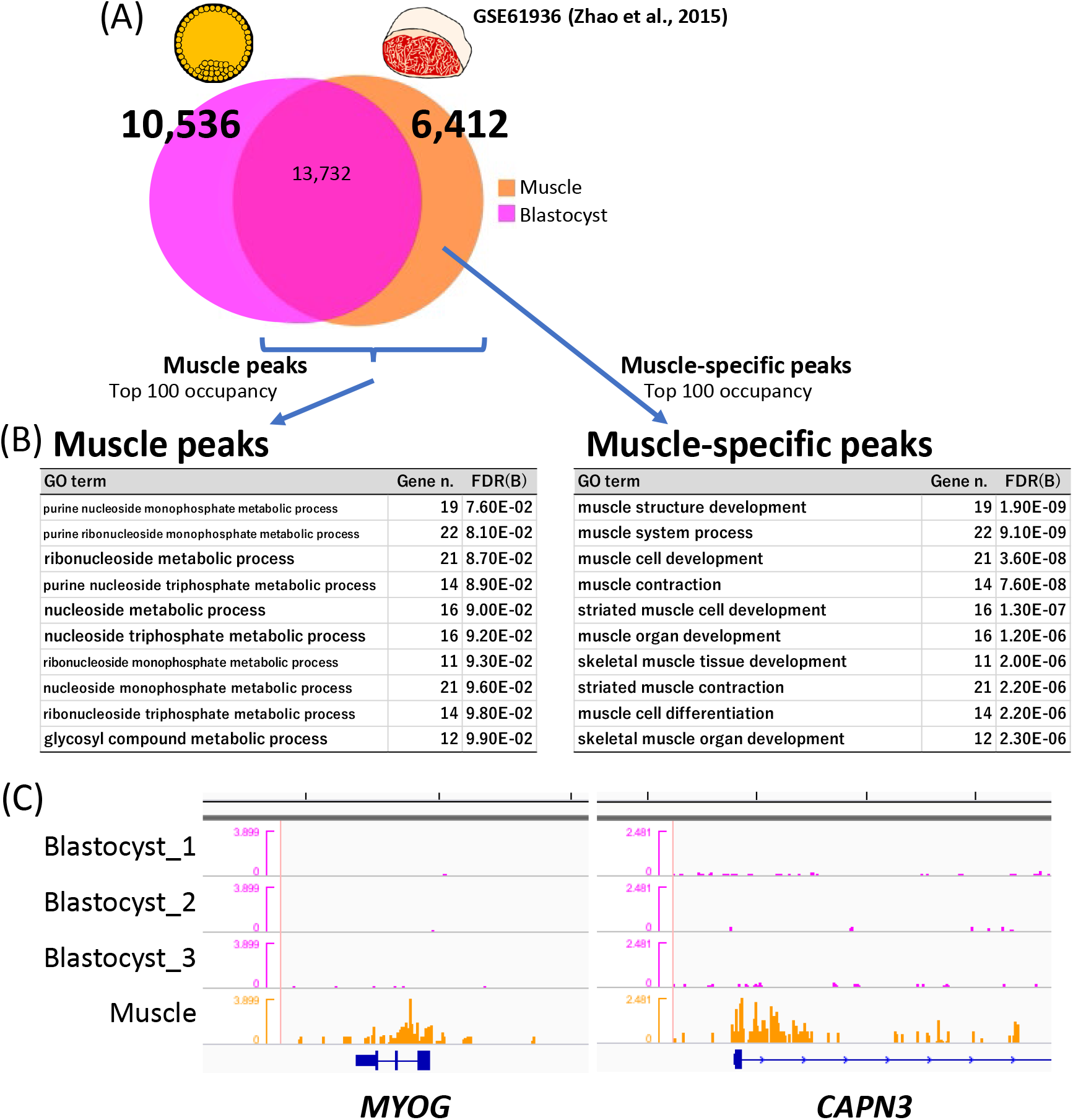
Characterization of blastocyst- and muscle-specific H3K4me3 modifications. (A) Processing of ChIP-peaks in muscle and blastocyst samples. The numbers show those of the peaks. The sum of common (intersect) and specific peak numbers is not identical to the original peak number in each sample because some peaks were separated into multiple peaks to represent intersects. (B) Top 10 significant GO terms for biological process enriched by the genes with the top 100 highest peak occupancy rates. Gene n. represents the numbers of related genes. FDR(B) indicates the Benjamin false discovery rate. (C) Examples of the H3K4me3 landscape for muscle-specific peaks (*MYOG* and *CAPN3*) around their TSSs. Five-kb graduations are shown in the top scale.

### 3.3 Characterization of common H3K4me3 modifications in blastocysts and liver

We next characterized the genes that harbored common H3K4me3 modifications in blastocyst and liver. Among the publicly available bovine H3K4me3 data from somatic tissues (liver and muscle), the liver data were relatively comparable with our blastocyst data in terms of available read numbers, average profiles of the gene groups of interest (Figs. 4A and 5A), and the enrichment profiles of representative positive regions (Fig. S2A). However, the muscle data did not meet these criteria (Fig. S2A and B) and we did not analyze these data further. At first, we focused on meat production-related genes given that meat production is one of the most important economic traits in beef cattle. We used the list of meat production-related genes produced by Williams et al. on the basis of biological roles, which might influence muscle development, structure, metabolism, or meat maturation (Williams, et al., 2009). The 504 listed genes were curated with RefSeq mRNA accession numbers and official gene symbols, which narrowed down the gene number to 438. Of these, 216 harbored H3K4me3 peaks within ±3,000 bp of TSSs both in blastocysts and liver, and 203 genes were recognized by the ngsplot program (Shen, et al., 2014) to calculate peak area around the TSSs. The processed gene list is shown in Table S3. Figure 4A shows the average profile plot of the 203 genes in the samples (three blastocyst and two liver samples). Given that the overall profiles were similar, we considered that the study-dependent bias in the peak area was negligible and the peak area could be compared among the samples. Therefore, the peak areas at these genes were subjected to PCA. As a result, PC1 well characterized the inter-tissue differences of H3K4me3 modifications, the levels of which are largely different between blastocysts and liver, even though they are common modifications (Fig. 4B and C). For example, *CROT* had lower H3K4me3 levels in blastocysts (autoscaled area [mean ± standard deviation], −0.68 ± 0.26) compared with liver (1.02 ± 0.62), whereas *ALDH5A1* exhibited the opposite pattern (0.71 ± 0.23 for blastocysts and −1.07 ± 0.14 for liver). The genes largely contributing to PC2 exhibited relatively large intra-tissue deviations; for example, *PPP3CA* exhibited −0.21 ± 1.01 in blastocysts and 0.32 ± 1.25 in liver (Fig. 4C).

**Figure 4.**
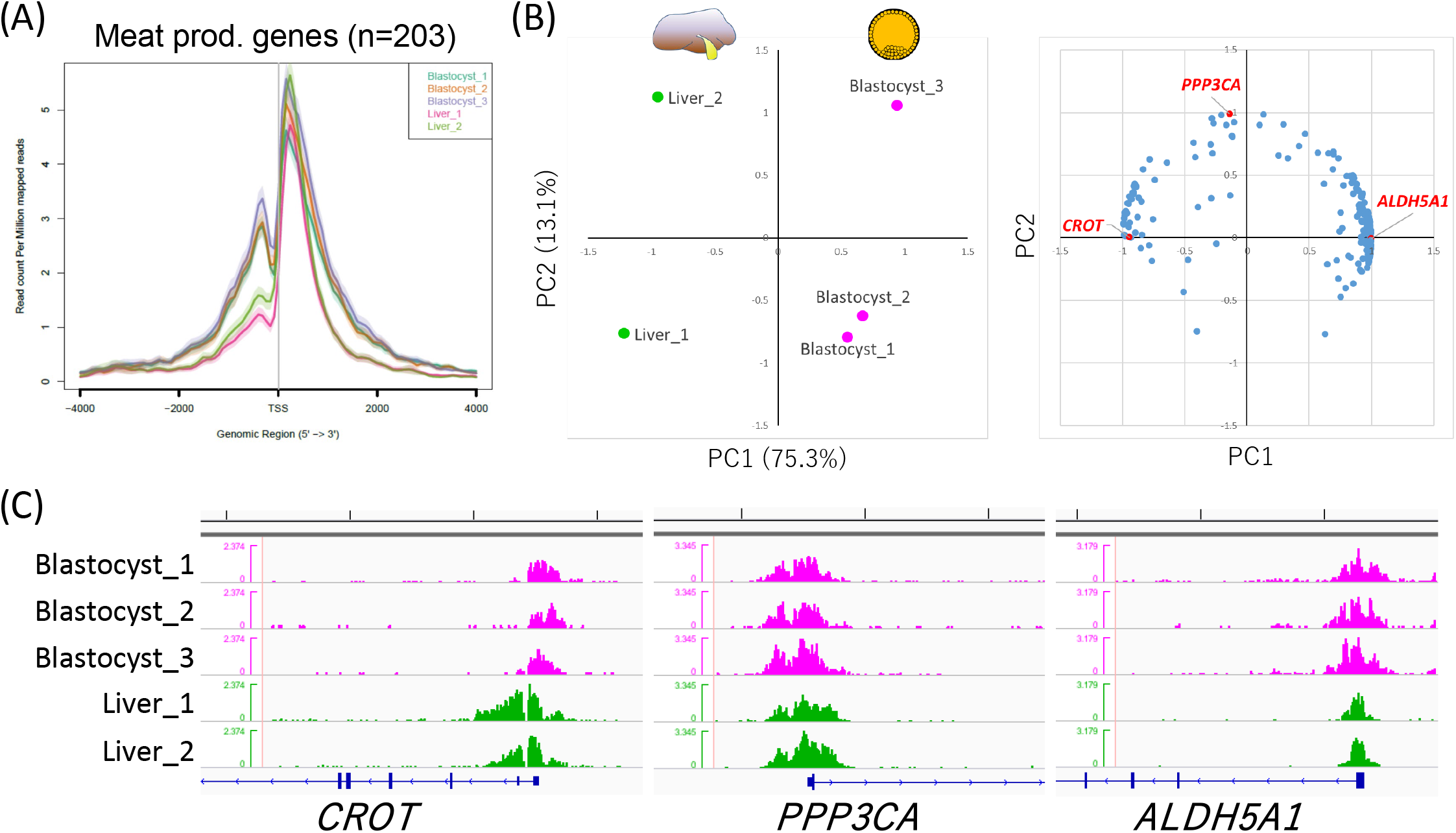
Characterization of H3K4me3 modifications common in blastocysts and liver in terms of meat production-related genes. (A) Average profile plot of the H3K4me3 signal around the TSSs of 203 meat production-related genes. (B) PCA of liver and blastocyst samples considering the peak areas around the TSSs of 203 meat production-related genes. Left and right panels show the principal component plot of all samples and loading plot of the 203 genes, respectively. The genes whose H3K4me3 landscapes are shown in (C) are highlighted in red. (C) H3K4me3 landscapes of *CROT, PPP3CA*, and *ALDH5A1* around their TSSs. Five-kb graduations are shown in the top scale.

**Figure 5.**
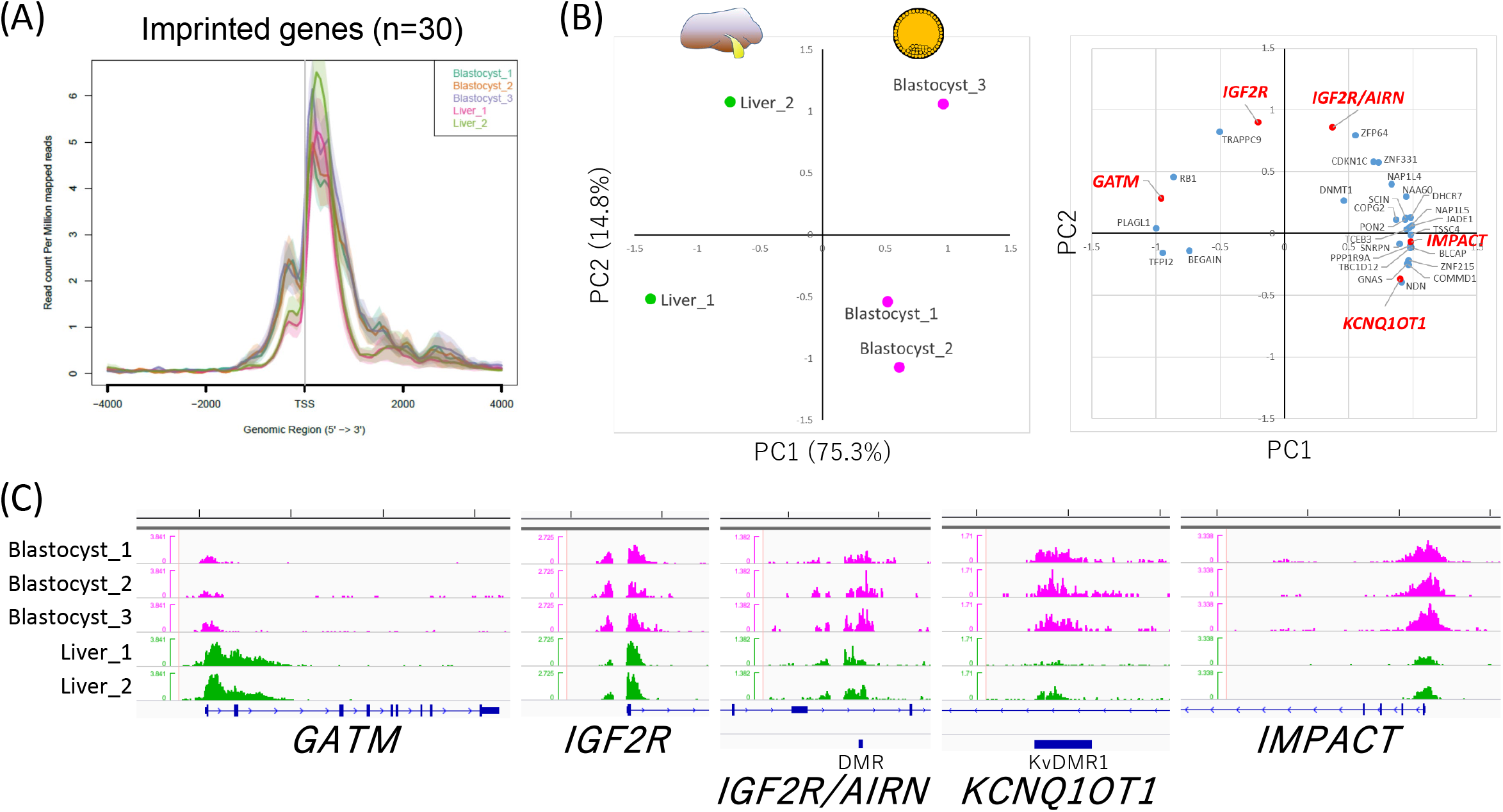
Characterization of H3K4me3 modifications common in blastocysts and liver in terms of imprinted genes. (A) Average profile plot of the H3K4me3 signal around the TSSs of 30 imprinted genes. (B) PCA of liver and blastocyst samples considering the peak areas around the 30 TSSs of the imprinted genes and two additional DMRs. Left and right panels show the principal component plot of all samples and loading plot of the loci examined, respectively. The genes whose H3K4me3 landscapes are shown in (C) are highlighted in red. (C) H3K4me3 landscapes of *GATM, IGF2R, IGF2R/AIRN* (DMR), *KCNQ1OT1* (DMR), and *IMPACT*. Five-kb graduations are shown in the top scale.

We then investigated H3K4me3 peaks that are common in blastocysts and liver in terms of imprinted genes given that these genes are profoundly involved in the etiology of fetal developmental disorders (Chen, et al., 2015) and genomic imprinting is also closely associated with the meat production traits of livestock (Neugebauer, et al., 2010; Okamoto, et al., 2019). The TSSs of 105 known imprinted genes listed in a previous bovine study (Chen, et al., 2015) and two differentially methylated regions (DMRs) related to bovine fetal overgrowth (KvDMR1 of *KCNQ1OT1* and *IGF2R/AIRN* DMR) were selected for investigation, and we found that 30 TSSs and the two DMRs had H3K4me3 peaks both in blastocysts and liver. The average profile plot of the 30 TSSs again exhibited overall similar profiles between the two different tissues (Fig. 5A). Meanwhile, we could also extract tissue-specific H3K4me3 modifications, as shown in Fig. S3. PCA of the common peaks in the two tissues extracted the inter- (PC1) and intra-tissue (PC2) differences represented by differential patterns of H3K4me3 modifications among the tissues and samples. The imprinted genes *IGF2R* (with *IGF2R/AIRN* DMR) and *KCNQ1OT1* (with KvDMR1), both of which are implicated in fetal overgrowth syndrome (Chen, et al., 2015), exhibited a relatively high contribution to intra-tissue differences (PC2).

## 4 Discussion

To our knowledge, only one study has assessed genome-wide histone modifications of bovine early embryos (Org, et al., 2019). That report was a pioneering study of the issue; however, due to its emphasis on an unconventional ChIP methodology to reduce sample cell numbers, H3K4me3 landscape that was produced was compromised by a lack of valley-like patterns around TSSs that correspond to nucleosome-free regions (Fig. S4) (Jiang and Pugh, 2009). The present ChIP-seq analysis of H3K4me3 successfully generated the typical landscape of epigenetic modifications around TSSs (Figs. 1D and Fig. S4). In addition, the clear enrichment of H3K4me3 modifications at TSSs and their correlation with the expression levels of the corresponding genes (Fig. 1) support the accuracy of our results. Therefore, the H3K4me3 profile produced in this study might be useful as a new reference for bovine early embryos.

We compared this high-resolution H3K4me3 profile of bovine blastocysts with those of somatic tissues (liver and muscle) to elucidate common and typical epigenetic modifications between early embryos and differentiated somatic tissues. Interestingly, the H3K4me3 peaks of blastocysts could be used like a “sieve” to extract somatic tissue-specific peaks in genes closely related to tissue-specific functions. Although the genes with the top 100 highest peak occupancy rates in blastocysts, liver, and muscle did not represent the specific functions of each tissue, their counterparts after subtracting genes with common peaks (i.e., “sieving”) clearly did (Figs. 2B and 3B and Table S2). These results suggest that high-throughput histone methylome data from early embryos are useful for sieving the methylome of other somatic tissues to characterize them in terms of the tissue-specific modifications that are related to their functions.

The common epigenetic modifications between early embryos and somatic tissues are important from the viewpoint of DOHaD, because they represent candidate modifications responsible for the epigenetic persistence-derived long-term consequences of early life conditions. In this study, the common peaks between blastocysts and liver were investigated with a focus on meat production-related genes. PCA of peak areas at TSSs well characterized inter-tissue differences (Fig. 4B and C); in other words, the genes highly contributing to PC1 represented H3K4me3 modifications that are largely different between blastocysts and liver, even though they are “common” modifications (e.g., *ALDH5A1* and *CROT*). On the other hand, PC2 might mirror intra-tissue differences rather than inter-tissue ones, for example, such as *PPP3CA* (Fig. 4C).

In addition to meat production-related genes, we also focused on imprinted genes. Genomic imprinting is an epigenetic phenomenon that compels a subset of genes to be monoallelically expressed in a parent-of-origin-dependent manner in mammals (Bartolomei and Ferguson-Smith, 2011). Appropriate monoallelic expression of imprinted genes is crucial for normal fetal growth, and environmental perturbation during early embryonic development, including the use of assisted reproductive technology, can induce a loss of imprinting of these genes, leading to abnormal fetal overgrowth syndrome (Chen, et al., 2015). Furthermore, genomic imprinting effects have been widely documented in the economic traits of livestock animals. For example, in beef cattle, several reports have described large relative imprinting variance (i.e., the proportion of total genetic variance attributable to imprinted genes) such as for fat score (24.8% for German Simmental bulls (Neugebauer, et al., 2010)) and beef marbling score (35.2% for Japanese Black bulls (Okamoto, et al., 2019)). Although DNA methylation has been characterized as a major code of genomic imprinting, recent studies reported DNA methylation-independent imprinting that is controlled by histone methylation (Inoue, et al., 2017). In addition, the global dysregulation of imprinted genes in assisted reproductive technology-induced fetal overgrowth is often independent of DNA methylome epimutations (Chen, et al., 2015; Chen, et al., 2017). These findings suggest the importance of histone methylation in the expression of imprinted genes. In the present study, we categorized H3K4me3 modifications on imprinted genes into tissue-specific (Fig. S3), common but relatively tissue-dependent (Fig. 5B, PC1 contributions), and common (Fig. 5B, PC2 contributions) modifications. Interestingly, the imprinted genes *IGF2R* and *KCNQ1OT1*, which are well documented in fetal overgrowth syndrome (Chen, et al., 2015), exhibited a relatively high contribution to intra-tissue differences (PC2), suggesting that these H3K4me3 modifications are common in early embryos and hepatic tissues and diverse among individuals and/or given conditions. Collectively, the appropriate controlling of these histone modifications in developmentally, and hence, economically important genes in the early life period might result in phenotypic changes to improve the welfare and production traits of farm animals. The effects of environmental conditions and the development-dependent changes on epigenetic modifications and the feasibility of controlling them remain targets for future research.

In conclusion, the present study produced a referential H3K4me3 landscape of bovine early embryos and revealed its common and typical features compared with adult tissues.

## Acknowledgments

The authors deeply thank the staff at the Kyoto-Meat-Market for allowing us access to bovine ovaries.

## Data Availability

The ChIP-seq datasets for bovine blastocysts have been deposited in the Gene Expression Omnibus of NCBI with accession number GSE161221.

## Author Contributions

M.I., S.I., and N.M. conceived the experiments and drafted the manuscript. M.I. performed bovine IVF and ChIP-seq library preparation and sequencing. M.I. and S.I. analyzed the ChIP-seq results. N.M. supported the experiments and analyses. All authors discussed the results and approved the manuscript.

