## Supplementary Information for "Comparative analysis of histone H3K4me3 modifications between early embryos and somatic tissues in cattle"

**Supplementary figure legends**

Figure S1. Correlation analysis of three biological replicates of ChIP-seq for H3K4me3 in bovine blastocysts. Scatterplots of three pairwise comparisons are shown with Pearson correlation coefficients. Ten kb bin size was used for drawing using deepTools (https://deeptools.readthedocs.io/en/develop/).

Figure S2. (A) H3K4me3 landscapes of the housekeeping genes *GAPDH* and *SDHA* derived from blastocyst, liver, and muscle ChIP-seq data. (B) Average profile plot of H3K4me3 signals around the TSSs of 165 meat production-related genes (left) and 20 imprinted genes (right), that harbor the modifications within ±3,000 bp of the TSSs both in blastocysts and muscle.

Figure S3. (A) Blastocyst- and (B) liver-specific H3K4me3 peaks around the TSSs of imprinted genes.

Figure S4. Comparison of the average profile plots of H3K4me3 signals around the genome-wide TSSs between the present and previous studies (1). Replicate 1 of the published data was used for plotting.

Table S1. Data generated by ChIP-seq analysis of bovine blastocysts using an anti-H3K4me3 antibody.

Table S2. Top 10 significantly enriched GO terms derived from liver- and blastocyst-peaks, respectively, without sieving.

Table S3. Meat production-related genes analyzed in this study.

1.	T. Org *et al.*, Genome-wide histone modification profiling of inner cell mass and trophectoderm of bovine blastocysts by RAT-ChIP. *PLoS One* **14**, e0225801 (2019).
