## Supplementary figures and images for "Comparative analysis of histone H3K4me3 modifications between early embryos and somatic tissues in cattle"

### Supplementary Fig

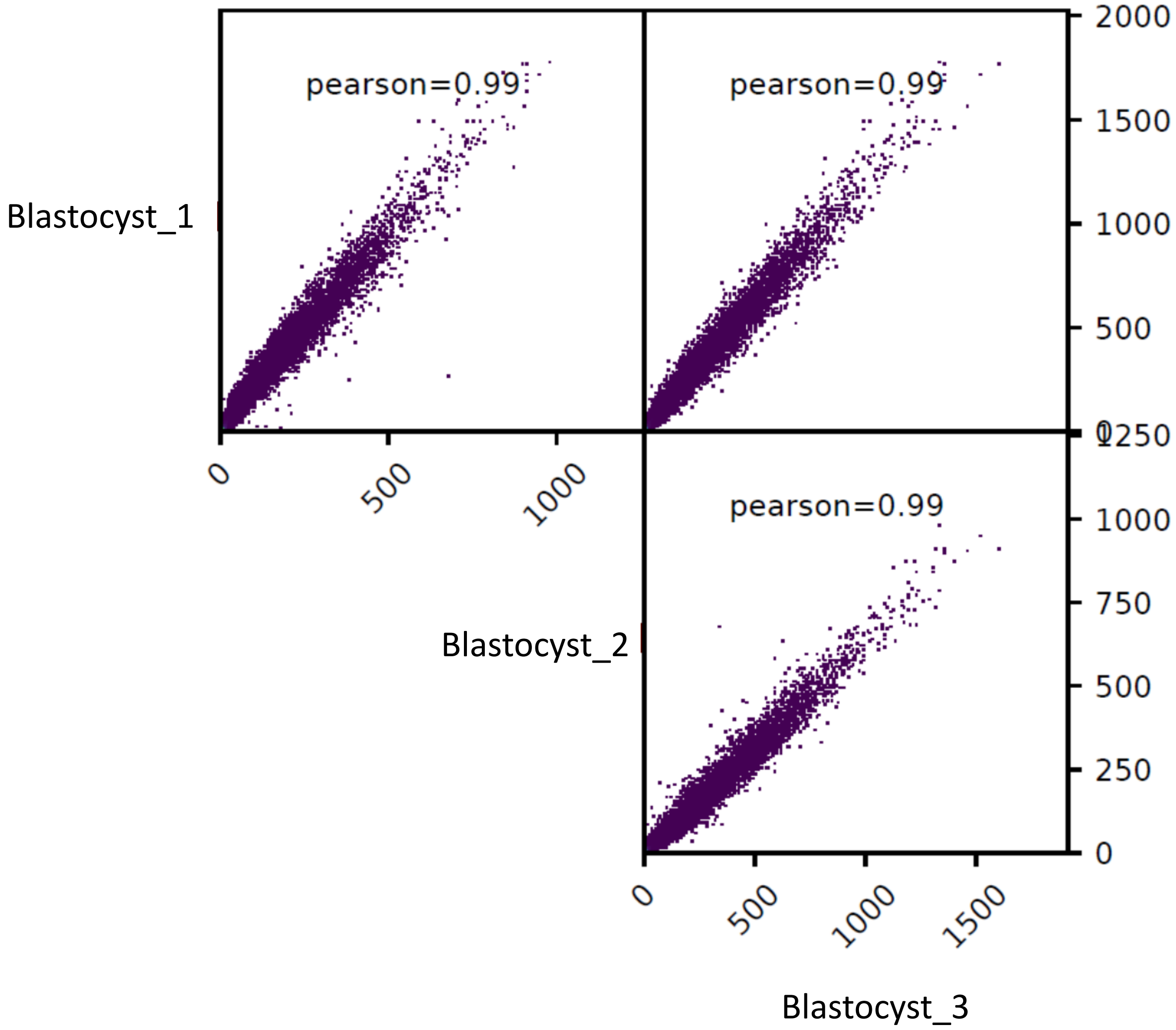

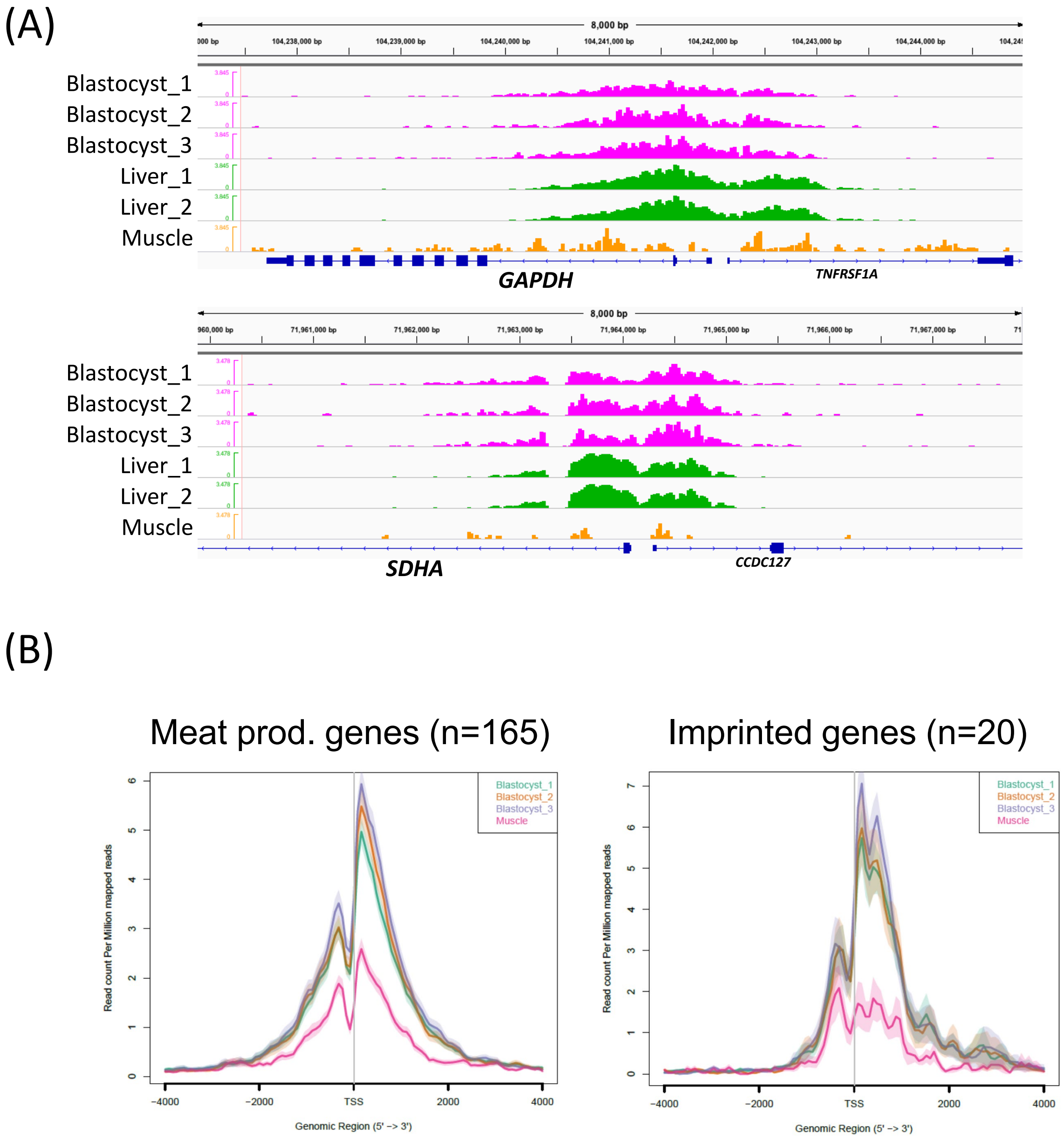

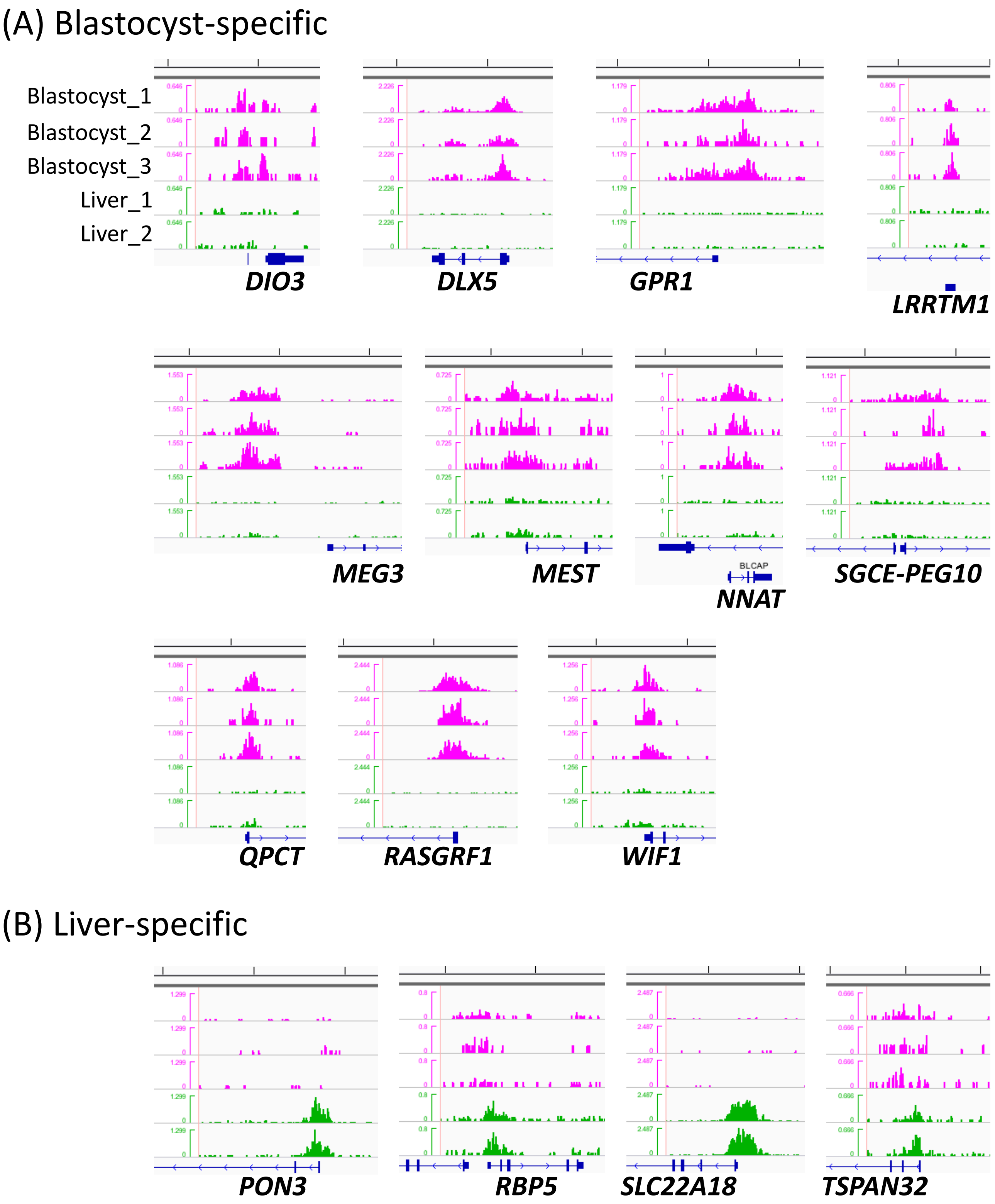

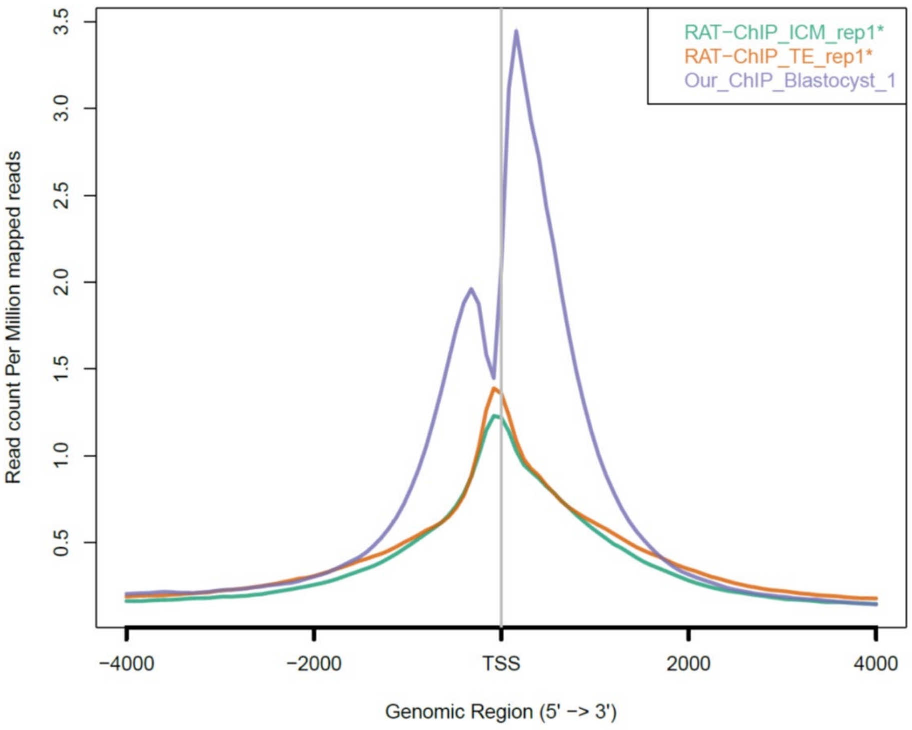
